# Viral fitness determines the magnitude of transcriptomic and epigenomic reprogramming of defense responses in plants

**DOI:** 10.1101/2019.12.26.888768

**Authors:** Régis L. Corrêa, Alejandro Sanz-Carbonell, Zala Kogej, Sebastian Y. Müller, Sara López-Gomollón, Gustavo Gómez, David C. Baulcombe, Santiago F. Elena

## Abstract

Although epigenetic factors may influence the expression of defense genes in plants, their role in antiviral responses and the impact of viral adaptation and evolution in shaping these interactions are still poorly explored. We used two isolates of turnip mosaic potyvirus (TuMV) with varying degrees of adaptation to *Arabidopsis thaliana* to address these issues. One of the isolates was experimentally evolved in the plant and presented increased load and virulence relative to the ancestral isolate. The magnitude of the transcriptomic responses were larger for the evolved isolate and indicated a role of innate immunity systems triggered by molecular patterns and effectors in the infection process. Several transposable elements (TEs) located in different chromatin contexts and epigenetic-related genes were also affected. Correspondingly, mutant plants having loss or gain of repressive marks were, respectively, more tolerant and susceptible to TuMV, with a more efficient response against the ancestral isolate. In wild-type plants both isolates induced similar levels of cytosine methylation changes, including in and around TEs and stress-related genes. Results collectively suggested that apart from RNA silencing and basal immunity systems, DNA methylation and histone modification pathways may also be required for mounting proper antiviral defenses in plants and that the effectiveness of this type of regulation strongly depends on the degree of viral adaptation to the host.

## INTRODUCTION

Biotic stress responses in plants can be triggered by the recognition of pathogens’ conserved motifs, proteins or RNA molecules. Pathogen-associated molecular patterns (PAMP) may be recognized by membrane receptors, triggering a general response referred as PAMP-triggered immunity (PTI) (Boutrot and Zipfel, 2017). A stronger defense is initiated when pathogen-specific proteins or other elicitors are recognized by resistance (R) proteins belonging to the NLR (intracellular nucleotide binding site, leucine-rich repeat containing receptor) family (Cui et al., 2015). The effector-triggered immunity (ETI) is linked to the induction of hypersensitive response (HR), restricting pathogen spread. Both PTI and ETI are associated with the production of hormones that may promote systemic resistance, inducing the production of resistance pathogenesis-related (PR) proteins, among others (Fu and Dong, 2013). Basal immunity systems are linked mainly to non-viral pathogens, but there is increasing evidence that they may also play a role against viruses (Teixeira et al., 2019).

RNA-based immunity systems are triggered by the recognition and degradation of double-stranded RNA (dsRNA) molecules. The mechanism is mostly associated in the defense against viruses (Wu et al., 2019). Viral dsRNAs are degraded by DICER-LIKE (DCL) proteins into small RNAs (sRNAs) that are loaded into ARGONAUTE (AGO) proteins and used as a guide to repress similar single-stranded RNAs. By using RNA-DEPENDENT RNA POLYMERASE (RDR) proteins to generate new dsRNAs from targets, the RNA silencing response can also be amplified (Borges and Martienssen, 2015). Viral dsRNAs can also feed into the PTI pathway (Niehl and Heinlein, 2019). Pathogens on the other hand, may evolve mechanisms to avoid or inactivate various steps of RNA silencing or PTI/ETI defenses, leading to Red Queen coevolutionary dynamics.

RNA-based defenses against viruses in plants are part of a broader and conserved system that includes processes that regulate gene expression and control transposable elements (TEs) by the addition of epigenetic marks to DNA or DNA-associated histone proteins (Borges and Martienssen, 2015). Most of the DNA methylation marks in eukaryotes are linked to cytosine, particularly those followed by guanine (CG). Non-CG methylation, including CHG and CHH (where H is any nucleotide, except G), however, is also observed. In plants, the symmetrical CG and CHG methylation are maintained by methyltransferases and the chromatin remodeling factor DECREASE IN DNA METHYLATION 1 (DDM1) during the replication process (Sigman and Slotkin, 2016). Signals for restoring asymmetrical CHH modifications, however, are lost and re-established after every cell division by a sRNA-guided complex. The mechanism known as RNA-directed DNA methylation (RdDM) is orchestrated by complexes containing two plant-specific RNA polymerases (Pol IV and V), the RNA silencing-related factors RDR2, DCL3 and AGO4 for sRNA generation and amplification and epigenetic factors, *e*.*g*., the methyltransferase DOMAINS REARRANGED METHYLASE 2 (DRM2) (Zhang et al., 2018).

RdDM mainly targets small and recently acquired TEs or the borders of long TEs in euchromatic regions (Sigman and Slotkin, 2016). The mechanism, therefore, establishes a heterochromatin-like environment within the euchromatin. Environmental stresses may pose a challenge for the maintenance of this chromatin border, as genes and TEs can mutually influence their expression under certain conditions (Negi et al., 2016). Changes in cytosine DNA methylation patterns due to stress have also been reported (Zhang et al., 2018). The impact of those epigenetic changes in gene expression settings are still elusive, especially for small and heterochromatin-poor genomes like the *Arabidopsis thaliana* one. The role of pathogens, and especially RNA viruses in DNA methylation responses also remains poorly explored. Contrary to passive abiotic stressors, pathogens can interact and manipulate host signaling pathways and therefore potentially exploit the intensity or types of epigenetic responses. In particular, fast-evolving RNA viruses may overcome host defenses by (*i*) quickly generating extremely diverse mutant swarms that contain escape variants that are not controlled by immunity (Andino and Domingo, 2015) or (*ii*) by encoding specific proteins that actively interact and block host defenses, being the viral suppressors of RNA silencing (VSR) relevant players in the context of this study (Wu et al., 2010).

We used *A. thaliana* ecotype Col-0 and *Turnip mosaic virus* (TuMV; genus *Potyvirus*, family *Potyviridae*; picorna-like superfamily) pathosystem as a model to explore those topics. TuMV is an economically relevant virus that infects cruciferous plants, including arabidopsis. Its compact positive single-stranded, polyadenylated RNA genome produces a single polyprotein that is processed into 10 major multifunctional proteins (Ivanov et al., 2014) plus an additional protein encoded in an alternative small ORF (Chung et al. 2008). To test whether viral evolution and adaptation changes the way viruses might interplay with host epigenetic regulation, two TuMV isolates with different fitness in arabidopsis were used. We show that epigenetic pathways have relevant roles in virus infectivity and that the responses are influenced by pathogen’s fitness in the host. We also find several virus-induced DNA methylation changes, but that their impact on transcriptional changes cannot be generalized. Overall, no major differences in the methylome exist between both viral isolates, however, the high-fitness TuMV isolate has a much stronger impact at the transcriptomic level.

## RESULTS

### Experimental evolution of TuMV in arabidopsis

Host-pathogen interactions in plants, as in other systems, are heavily regulated by a coevolutionary arms race of defense and counter-defense mechanisms. To check the impact of virus-acquired adaptations on host’s transcriptome and methylome responses, a calla lily isolate of TuMV (Chen et al., 2003), that had been propagated in *Nicotiana benthamiana* plants, was experimentally evolved by serial passages in arabidopsis plants (Figure 1A). By repeatedly challenging the virus population with the novel host, we expected to evolve a TuMV isolate better adapted to this particular host than the original isolate, which was naïve to the plant.

**Figure 1.**
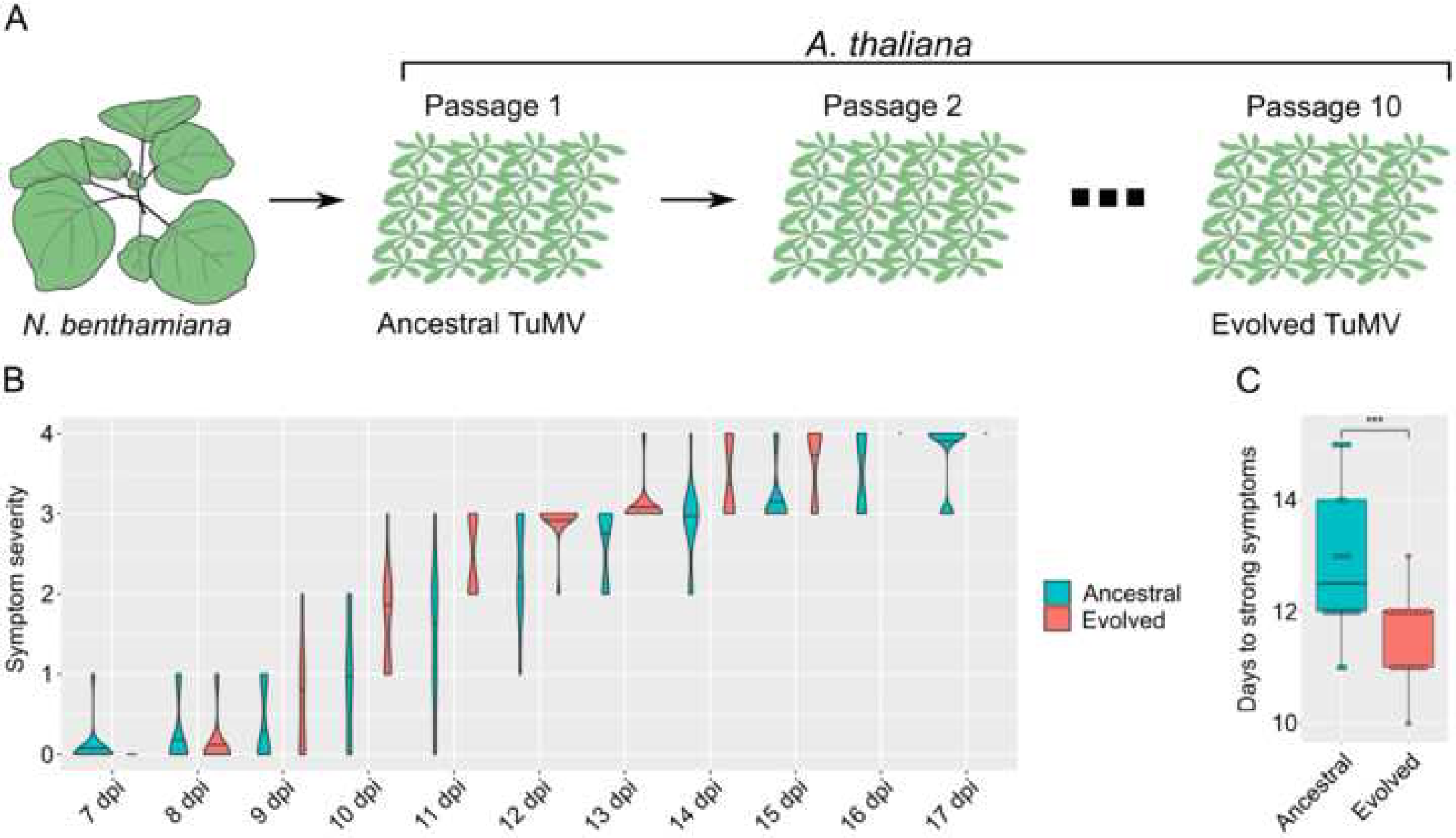
Experimental evolution of TuMV in arabidopsis. (A) A TuMV stock originally obtained from calla lily and subsequently maintained in *N. benthamiana* plants was used as a source of virus for an evolution experiment by serial passages in batches of arabidopsis wild-type plants. The first and 10^th^ passages were the ancestral and evolved isolates used in all experiments, respectively. (B) Symptom severity associated with ancestral and evolved TuMV isolates from 7 to 17 dpi, according to the scale defined in Figure S1A. Violin plots represent the symptoms severity level of each of the 20 plants infected with the different isolates. Lines represent the median severity value in each time-point. (C) Number of days each plant (dot) took to reach strong symptoms (symptom level 3, according to the scale provided in Figure S1A) after TuMV inoculation. Student two-samples *t* tests, \*\*\**P* < 0.001; \*\**P* < 0.01; NS., not significant.

When similar amounts of inoculum were used, the onset of early symptoms in the upper systemic leaves started ∼7 days post-inoculation (dpi), irrespective of the isolate used (Figure 1B). However, plants infected with the evolved virus progressed into strong symptoms faster than the ancestral-infected ones (Figure 1B and 1C). The largest symptom differences between the two viruses were observed 10 - 12 dpi (Figure 1B); most evolved virus-infected plants developed clear and strong leaf yellowing and stunning symptoms, while the ancestral virus-infected ones were still displaying light symptoms or remained symptomless (Figure 1B and S1A). The observed difference in symptoms was paralleled with viral load. At early infection stages (2 and 5 dpi), before symptoms appearance, there was no significant difference in the levels of TuMV accumulation (Figure S1B; 2-samples *t*-tests *P* ≥ 0.1620). However, after 12 dpi, when symptoms were clearly distinct between isolates, the load of the evolved virus was significantly higher than the ancestral virus in systemically-infected leaves (Figure S1B; 2 samples *t*-test *P* = 0.0013). In agreement, the evolved virus killed the plants significantly faster than the ancestral virus (Figure S1C; Kaplan-Meier survival analysis: *P* = 0.0003).

The genomes of the ancestral and evolved isolates were compared by variant calling through Illumina polyA-purified RNA sequencing (mRNA-seq) reads. Two single-nucleotide polymorphisms (SNPs) in the evolved isolate, leading to amino acid substitutions L107F and D110N, were observed (Figure S1D and S1E). Both substitutions affected the genome-linked viral protein VPg, which is a multifunctional protein involved in viral replication, genome stabilization, translation, and suppression of RNA silencing-based defenses (Cheng and Wang, 2017; Ivanov et al., 2014). These amino acids are located at the end of the third predicted α-helix (Figure S1D), in a region required for the VPg self-interaction and in close proximity to regions important for its interaction with the VSR protein HC-Pro and the host translation initiation factor eIF4E in related viruses (Roudet-Tavert et al., 2007; Yambao et al., 2003). Collectively, the development of symptoms, virus accumulation and molecular data indicated that the evolution experiment was effective in producing a TuMV isolate that is more virulent and better adapted to arabidopsis.

### Transcriptomic responses to TuMV infection

The magnitude and nature of plant transcriptomic responses to infection depend on the fitness of the particular potyviral strain being inoculated (Agudelo-Romero et al., 2008; Cervera et al., 2018; Hillung et al., 2016), including one study comparing two other TuMV strains (Sánchez et al., 2015). To confirm this observation in our particular pathosystem, arabidopsis plants were inoculated with equivalent amounts of transcripts from the ancestral or evolved TuMV isolates or mock-inoculated, and RNAs extracted from systemic leaves before symptom appearance (5 dpi) and late infection (12 dpi). In addition, a sample was taken 2 dpi (early infection) for the evolved virus. Transcriptomes of three biological replicates (plants) from each condition (mock-inoculated, ancestral virus- and evolved virus-infected) were assessed with stranded mRNA-seq. The vast majority of the reads in the infected plants mapped to the arabidopsis genome (Figure S2A), allowing the detection of differentially expressed genes (DEGs) in all time-points.

When compared to mock-inoculated samples, the number of DEGs was larger in the response against the evolved virus in all time-points analyzed. The number of DEGs for the evolved virus at 2 and 5 dpi was about three and seven times higher than for the ancestral at 5 dpi, respectively (Figure 2A and Table S1), indicating that the evolved virus elicited stronger responses at 2 dpi than the ancestral at 5 dpi. As infection progressed, responses between isolates tended to equalize, although total number of DEGs were still ∼1.5 higher for the adapted virus at 12 dpi (Figure 2A and S2B). A total of 18 genes were regulated due to the infection in all time-points for both viruses (Figure S2B), including eight known stress-responsive genes (Table S2).

**Figure 2.**
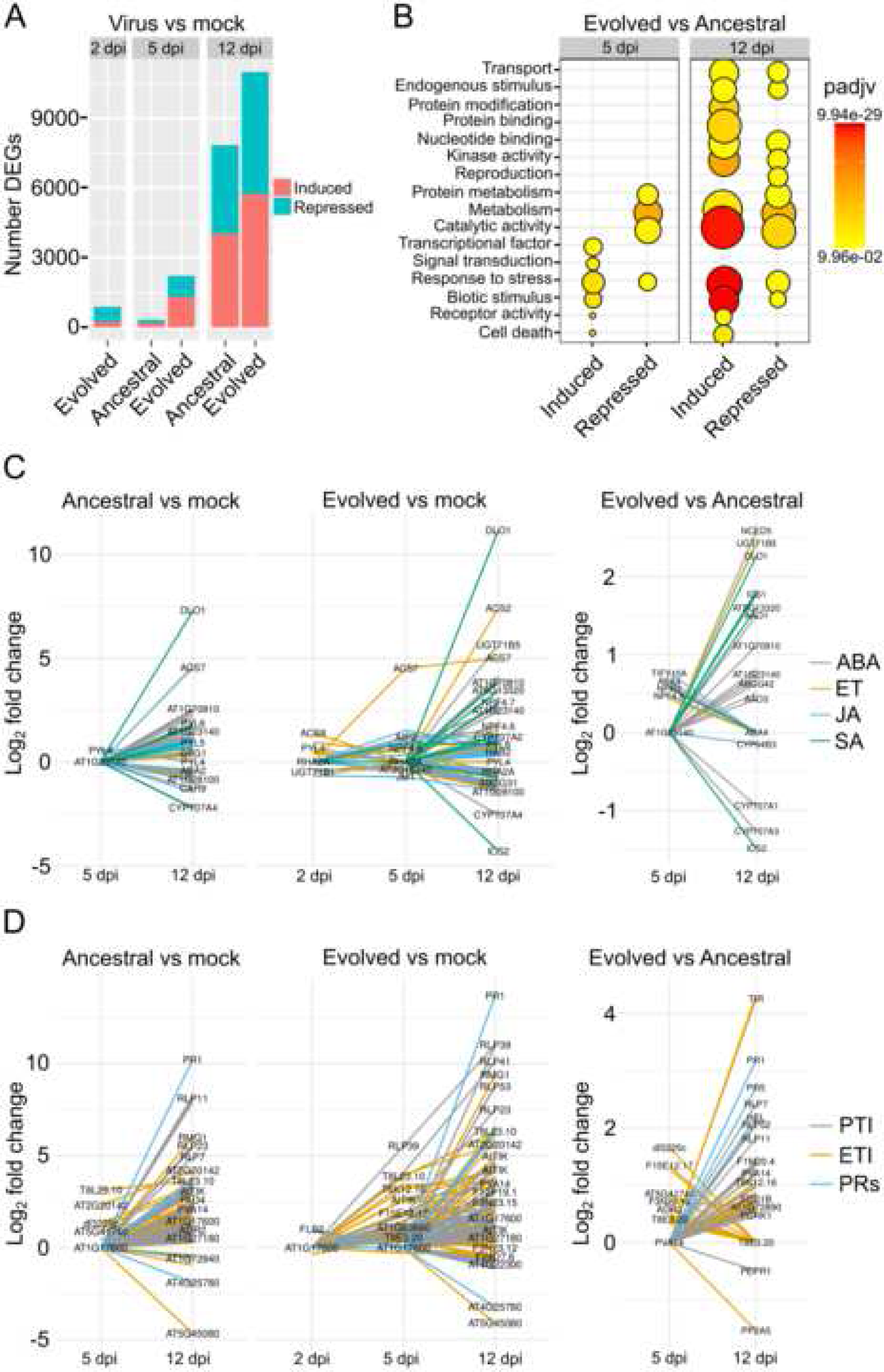
Transcriptome responses to TuMV. (A) Number of DEGs obtained by DESeq2 analysis for each TuMV infection condition (adjusted *P* < 0.05). In each time-point, three biological replicates infected with either the ancestral or evolved TuMV isolate were compared to mock-inoculated ones. (B) Gene ontology analysis (plant GOSlim) for DEGs between the evolved and ancestral TuMV isolates. For highlighting the differences between the isolates, TuMV ancestral- and evolved-infected samples were used as control and treatment in the DESeq2 analysis, respectively. Circle size represents level of enrichment and color heat maps indicate adjusted *P* values (padjv). (C) Transcriptional profiles (log_2_ fold change) of selected phytohormone genes, including abscisic acid (ABA), ethylene (ET), jasmonic acid (JA) and salicylic acid (SA). In the left and central panels both virus isolates were compared against mocks. In the right panel, the evolved isolate was directly compared against the ancestral one. (D) Transcriptional profiles (log_2_ fold change) of selected innate immunity genes, including PAMP-triggered immunity (PTI), effector-triggered immunity (ETI) and pathogenesis-related (PR) genes. As above, samples from evolved isolate-infected plants were directly compared against the ancestral isolate-infected plants in the right panel. dpi: days post-inoculation.

Responses against the ancestral virus at 5 dpi were characterized by an enrichment in genes associated with biotic and abiotic stresses and repression of metabolic and biosynthetic processes (Figure S2C). The core of defense-related genes associated with general stress-responses, though, were only observed at 12 dpi for plants infected with the ancestral isolate. Responses to the evolved virus, on the other hand, were much faster. At 2 dpi, typical shut-down of general metabolism and photosynthesis was already observed (Figure S2C). All major classes of regulated genes observed only at 12 dpi for the ancestral virus were already enriched against the evolved one at 5 dpi. At 12 dpi, those classes were enhanced, with the additional repression of ribosome constituents (Figure S2C). To highlight the difference between the viral isolates, the direct comparison of the transcriptomes from plants infected with the ancestral and evolved strains was performed. This analysis evidenced the stronger perturbation of the overall physiological homeostasis by the evolved isolate, including the induction of genes related to general and biotic stresses and transcriptional factors (Figure 2B). Suppression of biotic stress genes in evolved-infected samples was also observed, a change that may possibly be to the advantage of the virus (Figure 2B).

A network analysis of the identified DEGs was performed using the Arabidopsis Comprehensive Knowledge Network (AtCKN) (Ramšak et al., 2018). Dynamic views of AtCKN’s cluster 40, enriched for several well characterized stress-responsive genes, are available in the Supplemental Files S1 (ancestral) and S2 (evolved). The analysis indicated that the evolutionary conserved WRKY transcriptional factors may play important roles in response triggering and dynamics. At 2 dpi, *WRKY70*, a known activator of salicylic acid (SA)-related defense genes and a repressor of jasmonic acid (JA)-ones (Li et al., 2004), was induced against the evolved virus (Figure S2D). At 5 dpi, both isolates induced the expression of *WRKY25* and several of its direct targets (Figure S2D, Table S1, Files S1 and S2). This gene is a known repressor of SA responses (Zheng et al., 2007). Its induction, together with other WRKY SA-counteractors (*WRKY26*/*33*/*38*/*62*) evidenced a possible SA-buffering mechanism at mid and late infection points (Kc et al., 2008; Zheng et al., 2006). Other transcriptional factor families also seem to have relevant roles during early responses (Figure S3A and Table S3). A cross-talk with other hormones was also observed in early and late infection phases, especially genes related to abscisic acid (ABA), ethylene (ET) and JA (Figure 2C, Table S3). Furthermore, several genes associated with both PTI and ETI systems were regulated against TuMV, though to a larger extent for the evolved virus, with PR1 having the highest fold change among them at 12 dpi (Figure 2D, Table S3). A pronounced induction of PTI and PRs were observed in response to the evolved isolate at 12 dpi when compared to the ancestral one, which were paralleled with a higher increase of SA genes in this time-point (Figures 2C and 2D). ETI-related genes seem to be more dynamically regulated when the isolates are compared, with the bulk difference taking place at 5 dpi, despite the high induction of some of them at 12 dpi (Figure 2D). Expression of representative genes (*PR1, WAK1, HSP70*, and *COR15a*) associated with biotic and abiotic stresses were confirmed by quantitative PCR (RT-qPCR) to validate observed mRNA-seq results (Figure S3B), confirming that, on average, expression of these genes was higher in plants infected with the evolved strain (*post hoc* pairwise comparisons with sequential Bonferroni correction *P* = 0.0046).

Viruses are targeted by RNA silencing defenses. Accordingly, the majority of the RNA silencing-related genes among the DEGs were induced (Figure 3A, Table S3). Most of the DNA methylation-related DEGs, on the other hand, were repressed against both isolates (Figure 3A, Table S3). The changed expression of *INCREASE IN BONSAI METHYLATION 1* (*IBM1*) and *REPRESSOR OF SILENCING 1* (*ROS1*), two genes known to act as methyl sensors (Lei et al., 2015; Rigal et al., 2012), was confirmed by RT-qPCR (Figure S3C). Interestingly, the average level of expression was significantly lower in plants infected with the evolved than with the ancestral virus (*post hoc* pairwise comparisons with sequential Bonferroni correction *P* < 0.0001). The overall responses therefore indicated that both DNA/histone layers of epigenetic regulation might be altered during virus infection.

**Figure 3.**
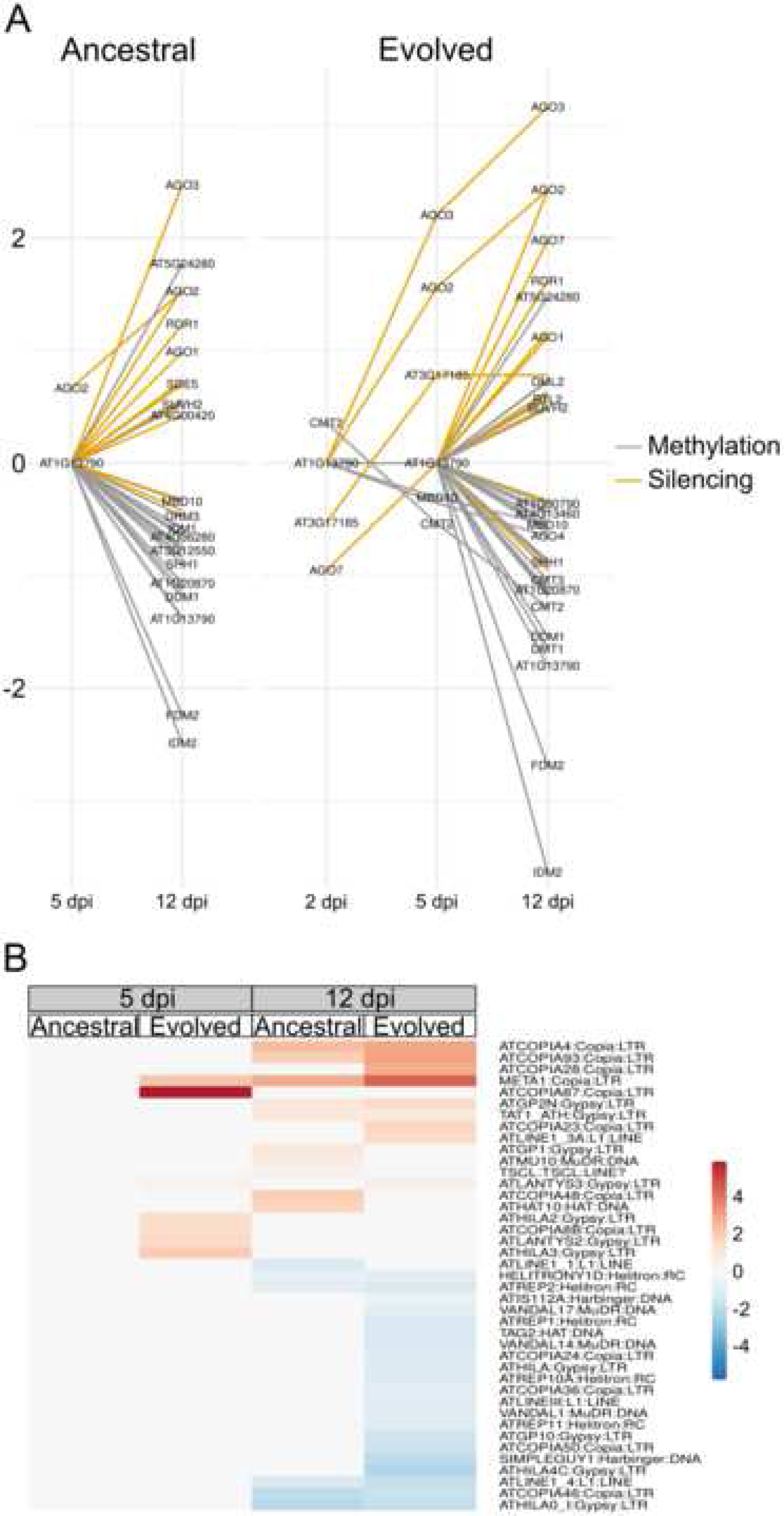
Transcriptional profiles of epigenetic-related selected genes and transposons. (A) Transcriptional profiles (log_2_ fold change) of selected RNA silencing (yellow lines) and DNA methylation genes (grey lines). (B) Heat map with fold changes of differentially expressed transposons (adjusted *P* < 0.05) obtained with the TEtranscripts tool. dpi: days post-inoculation.

Since several epigenetic pathways have TEs as targets, the expression of TE families was checked with TEtranscripts, a tool developed to handle reads mapping to repetitive sequences (Jin et al., 2015). At 5 dpi, seven TEs belonging to the Gypsy and Copia families, usually concentrated in centromeric and pericentromeric regions (Underwood et al., 2017), respectively, were induced against the evolved isolate (Figure 3B, Table S4). One of them, the Gypsy ATHILA2, is enriched in the centromere core that is transiently regulated by temperature shifts and viral infections (Diezma-Navas et al., 2019; Tittel-Elmer et al., 2010). At 12 dpi, however, both induction and repression of TE families was observed (Figure 3B, Table S4). The induced elements were again mostly from Gypsy and Copia families, including AtCOPIA93/Evadé, a TE that is induced against bacterial and viral infections (Diezma-Navas et al., 2019; Zervudacki et al., 2018). The repressed TEs at 12 dpi, on the other hand, included the Helitron, Harbinger and Mutator (MuDR) families that are usually located close to genes (Figure 3B, Table S4). The misregulation of several DNA methylation and histone modification genes and TEs located in different genomic contexts further suggested that epigenetic factors may play a role during the infection process.

### Effects of epigenetic-related genes in arabidopsis response to TuMV infection

So far, we have presented evidence suggesting that epigenetic factors may play a role during TuMV infection. To directly test this possibility, arabidopsis mutant genotypes having compromised or enhanced DNA/histone methylation were challenged against the two TuMV isolates. Disease severity was checked by scoring the number of days each plant took to reach strong leaf yellowing symptoms. All tested RdDM mutants, involved mainly in the regulation of small TEs located within euchromatic environments, were more resistant to TuMV than wild-type plants; though they were significantly more resistant against the ancestral isolate (Figure 4A; *post hoc* pairwise comparisons with sequential Bonferroni correction *P* < 0.0001). Among the challenged RdDM genotypes, *ago4* and *rdr2* were the most and least resistant ones, respectively, while *poliv, polv* and double *drm1 drm2* presented intermediate values. Strong resistance, especially for the ancestral virus, was also observed in *ddm1* mutants, lacking a master regulator of TEs (Figure 4B; *post hoc* pairwise comparisons with sequential Bonferroni correction *P* ≤ 0.0003).

**Figure 4.**
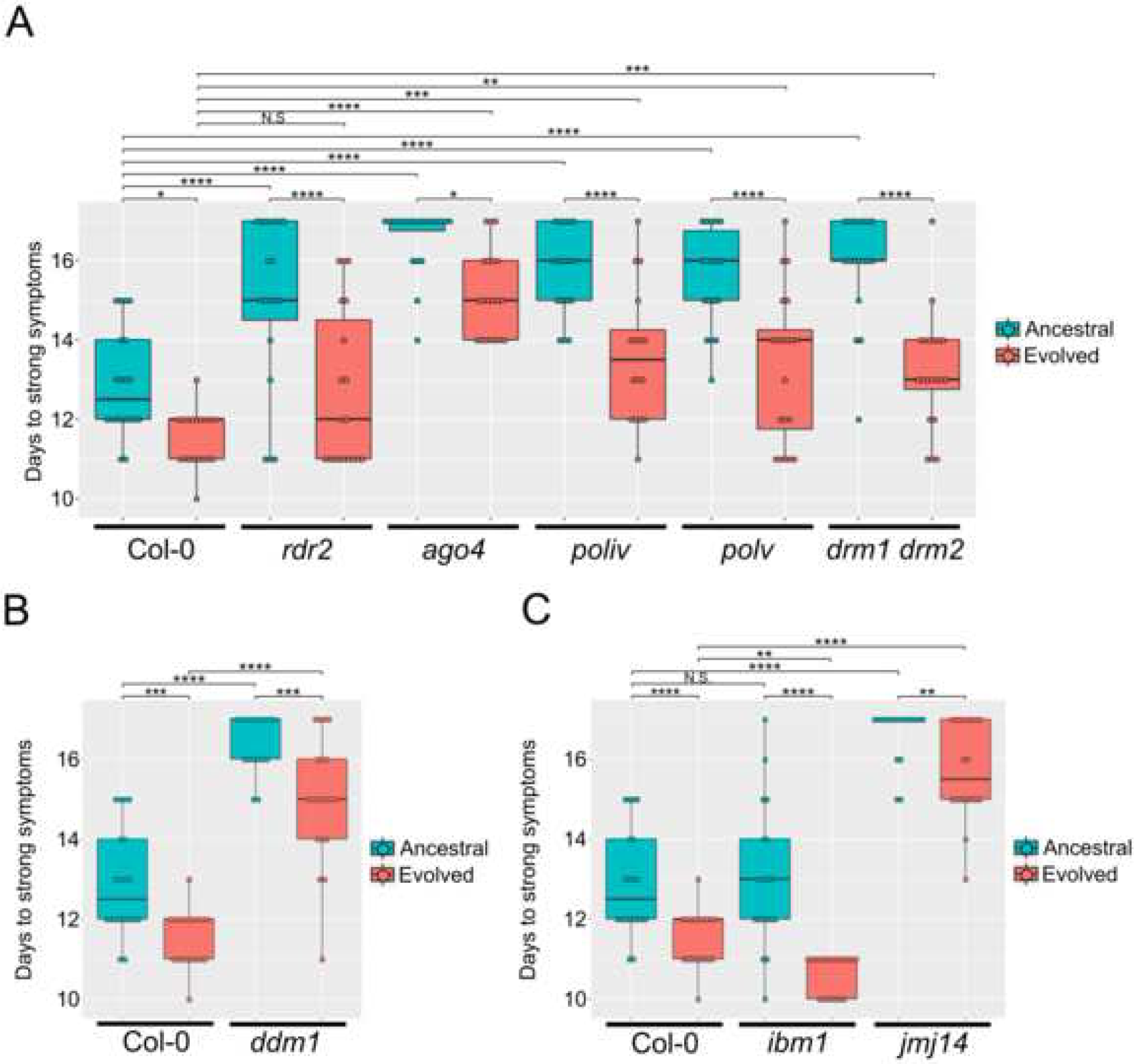
TuMV infection in epigenetic mutants. Number of days each plant (dot) took to reach strong symptoms after TuMV inoculation in epigenetic mutants, compared to Col-0 wild-type plants. (A) Panel of selected RdDM mutants. (B) Chromatin remodeler *ddm1* mutant. (C) Histone modification mutants. In all panels, the variable days to strong symptoms was fitted to generalized linear mixed models (GLMM) with plant (as indicated in the corresponding abscissa axes) and virus genotypes (ancestral and evolved) as orthogonal factors; a Normal probability distribution and an identity-link function were assumed. *Post hoc* pairwise comparisons with sequential Bonferroni correction tests were performed; \*\*\*\**P* < 0.001; \*\*\**P* < 0.01; \*\**P* < 0.05; \**P* < 0.1; NS., not significant.

Histone modification mutants, however, had opposite effects depending on the altered pathway. Compared to wild-type plants, *ibm1* mutants were significantly more susceptible to the evolved isolate (Figure 4C; *post hoc* pairwise comparisons with sequential Bonferroni correction *P* < 0.0001), but not against the ancestral one (Figure 4C; *post hoc* pairwise comparisons with sequential Bonferroni correction *P* = 0.7778). IBM1 is a histone demethylase that removes TE-associated H3K9 marks from genes, therefore reinforcing euchromatin/heterochromatin borders (Saze et al., 2008). On the other hand, inoculation of both isolates in mutants of the gene *JUMONJI 14* (*JMJ14*), rendered plants more resistant to the virus (Figure 4C; *post hoc* pairwise comparisons with sequential Bonferroni correction *P* < 0.0001). JMJ14 is also a histone demethylase, but removes H3K4 methylation marks, a modification usually associated with gene activation (Lu et al., 2010; Searle et al., 2010).

Infection results in the mutant backgrounds therefore indicated that infectivity and development of symptoms severity may be correlated to altered chromatin states. Mutants defective in heterochromatin formation (RdDM mutants, *ddm1* and *jmj14*) are more tolerant to TuMV infection, whilst the one with reduced euchromatin (*ibm1*) was more susceptible. The experiments also support the transcriptome findings that epigenetic factors may be required for virus defense mechanisms in plants.

### Virus-induced DNA methylation changes

Since several genes related to cytosine DNA methylation influenced TuMV infectivity, we asked whether this type of epigenetic modification is altered during the infection process in wild-type plants. Whole-genome bisulfite sequencing methylome libraries were constructed and Illumina-sequenced (WGBS-seq) for ancestral and evolved virus-infected plants at three time points: 2, 5 and 12 dpi. DNA material came from the same samples used for the transcriptome analysis. The observed differentially methylated regions (DMRs, in 100 bp tiles) were analyzed separately for the three cytosine methylation contexts (CpG, CHG and CHH) and divided into hypermethylated and hypomethylated, for gain or loss of methylation in comparison to mock-inoculated control plants, respectively (Table S5). In contrast to the transcriptome data, the numbers of DMRs induced by TuMV were relatively even between the ancestral and evolved isolates along the time-course (Figure S4A). The exception was for CHG DMRs at 12 dpi, that were clearly more pronounced in evolved virus-infected plants, with ca. twice of them hypermethylated. DMRs in the CpG context were in general more numerous during TuMV infection than in the other contexts (Figure S4A). CHG and CHH DMRs, however, had a marked increase at 12 dpi (Figure S4A).

Most of the observed CpG DMRs were mapped within protein-coding genes (Figure 5A and 5B). However, DMRs in CpG context proximal to transcriptional start sites (TSS) were also observed and, to a lesser extent, within TEs (Figure 5A and 5B). CHG and CHH DMRs, as expected, were enriched in TEs, with increased numbers in later infection times (Figure 5A and 5B). Plants infected with the evolved virus had about 2-fold more CHG DMRs in TEs at 12 dpi, corresponding to the bulk methylation difference in this context between the isolates (Figure 5A, 5B and S4A). In agreement with transcriptional profiles, DMRs from all three contexts were found in TE families located throughout the genome, with Gypsy, MuDR and Copia the most frequent ones (Figure S4B). While CpG DMRs in TEs tented to have similar amounts of hyper- or hypo-methylation irrespective of the time point, non-CpG DMRs in those elements had a clear tendency for hypermethylation at 12 dpi (Figure 5A, 5B and S4B). Methylome profiles identified therefore the existence of DMRs in both TEs and genes during TuMV infection, suggesting a possible mutual regulation between them.

**Figure 5.**
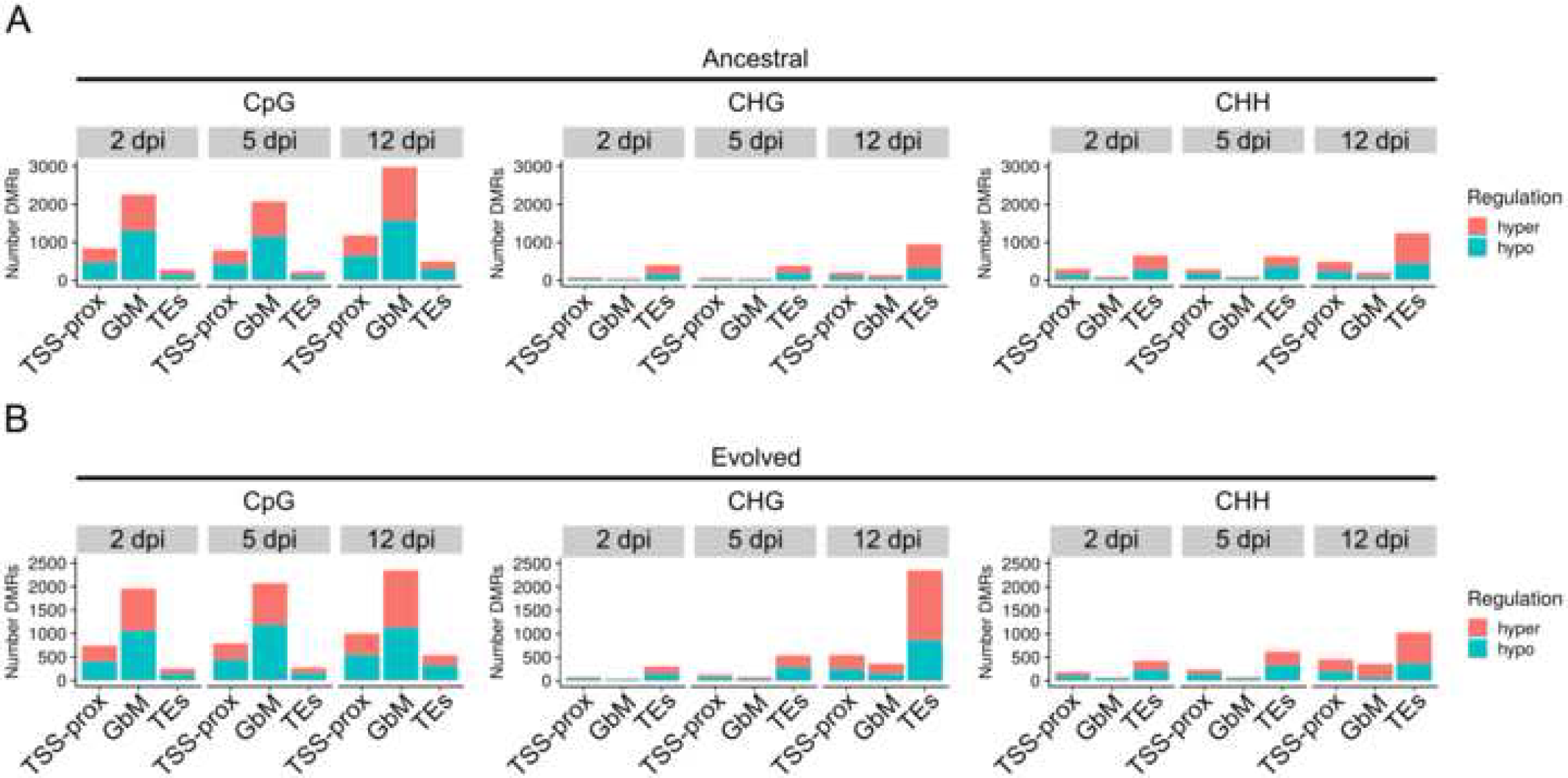
Whole-genome bisulfite sequencing (WGBS) of infected wild-type plants. Number of hyper- or hypo-methylated differentially methylated regions (DMRs) in the three cytosine contexts (CpG, CHG and CHH) proximal to transcriptional start sites (TSS-prox), inside genes (GbM) or TEs. (A) Ancestral TuMV-infected plants. (B) Evolved TuMV-infected plants.

### Impact of virus-derived methylation changes on the transcriptome

Since methylation of promoters is usually associated with changes in gene expression, the impact of TSS-proximal DMRs in the expression of protein-coding genes was assessed. DMRs in the CpG context were the most abundant ones in the region comprising 2 kb upstream and 200 bp downstream of protein-coding genes’ TSS, followed by CHG and CHH DMRs. If TSS-proximal methylation has a role in gene expression control, a negative correlation between them would be expected. However, most genes having TSS-proximal DMRs were not regulated by the infection at any time-point, regardless of the context (Figure 6A). Cases of negative correlation between TSS-proximal methylation and expression, though, were observed, especially in the CpG context at 12 dpi (Figure 6A, Table S6). The observed correlations were mainly linked to TSS-proximal hypermethylation and repression of gene expression, although few cases in the opposite direction were also observed (Table S6). These genes were classified according to functional categories. Genes related to RNA metabolism (biosynthesis and processing) and protein metabolism (modification and translocation) were the most predominant ones (Figure 6B). Genes related to amino acid, carbohydrate, coenzyme, lipid, nucleotide, and secondary metabolism were also enriched. Despite being one of the most responsive in the transcriptome, few stress-related genes had negative correlation with DMRs (Figure 6B, Table S6).

**Figure 6.**
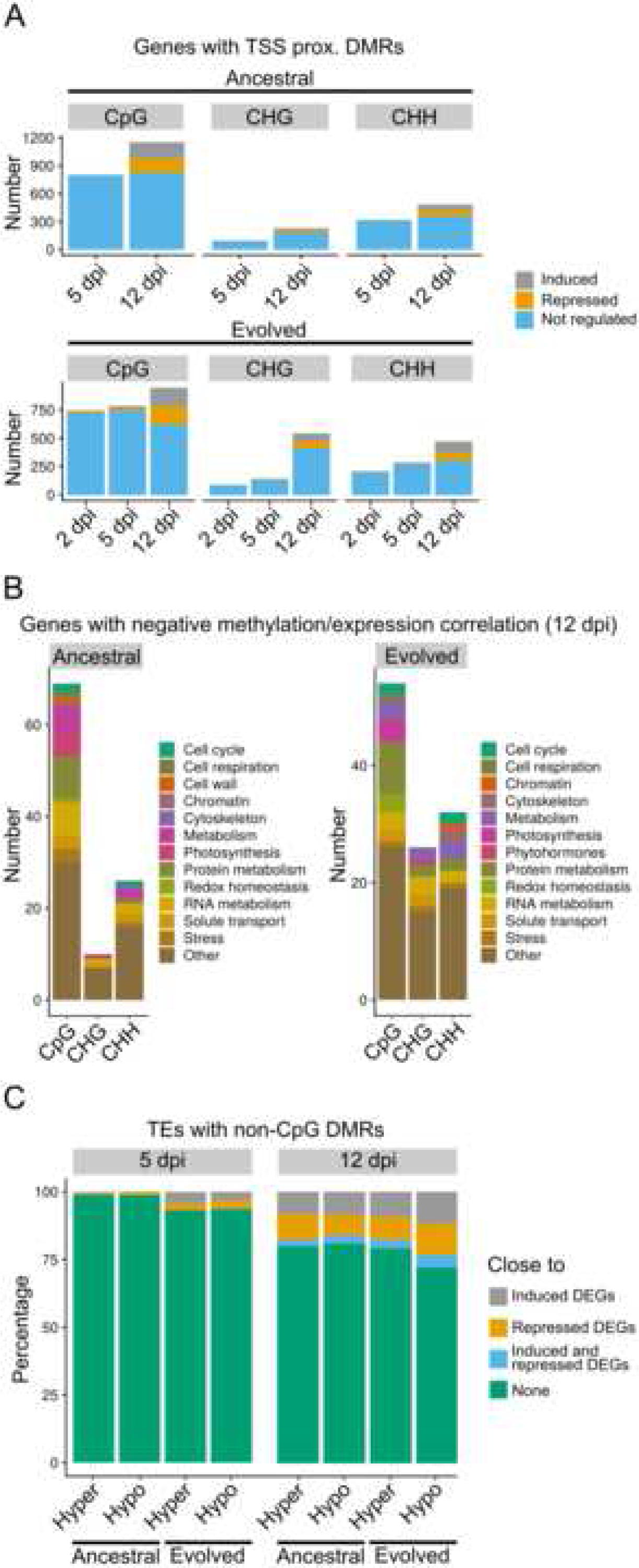
Correlation between TuMV-induced methylation and expression. (A) Number of genes having differentially methylated regions (DMRs) proximal to transcriptional start sites (TSS-prox) that were found to be regulated at the transcriptional level. (B) Functional characterization based on MapMan bins of genes having negative correlation between TSS-prox methylation and expression. (C) Percentage of TEs with hyper- or hypo-methylated non-CpG DMRs that are close (up to 10 kb) or far from DEGs.

Since CHG and CHH are the major transposon-associated methylation marks and that variation in their patterns can influence the expression of nearby genes (Sigman and Slotkin, 2016), we also sought for cases where elements with non-CpG DMRs were close to virus-induced DEGs. At 5 dpi, the vast majority of methyl-regulated TEs were further than 10 kb from DEGs, indicating that either their regulation did not influence expression of nearby genes or that they were located outside of gene-rich areas (Figure 6C). A larger number of regulated TEs close to DEGs was observed at 12 dpi, although elements far from DEGs were still the predominant type (Figure 6C). In both time points, there was no clear general correlation between the state of TE regulation (hyper- or hypo-methylated) and expression direction (induced or repressed) of nearby genes, probably reflecting the dynamic changes in their control along the infection time course. At 12, about 80 DEGs, including PTI- and ETI-related genes, were found to be close to regulated TEs in both isolates (Table S7). There were also isolate-specific cases: about 150 DEGs were close to regulated TEs in the ancestral and other 300 in the evolved TuMV. Abiotic and biotic stress-related genes were also found among the isolate-specific TE-close DEGs (Table S7).

## DISCUSSION

In this study we have used different approaches to evaluate the impact of epigenetic factors in triggering stress responses against viruses in plants. Infection experiments in epigenetic-deficient mutants indicated that RdDM factors, including AGO4, RDR2, POLIV, POLV and DMR1/2, the chromatin remodeler DDM1 and the histone modification proteins IBM1 and JMJ14 can control responses against TuMV infection (Figure 4). RdDM-, DDM1- and JMJ14-deficient plants showed resistance against the virus, while *ibm1* mutants were more susceptible. This agrees with experiments performed in inflorescence of other arabidopsis epigenetic mutants (*drm1 drm2, drm1, drm2, cmt3*, and *ros1*) infected with a tobravirus (Diezma-Navas et al., 2019). Other studies have also associated loss of DNA methylation factors with increased resistance against non-viral biotrophic pathogens, but susceptibility to necrotrophic ones (Dowen et al., 2012; Le et al., 2014; López et al., 2011; López Sánchez et al., 2016; Luna et al., 2012; Yu et al., 2013). Biotrophic pathogens are thought to be targeted mainly by SA-mediated defenses and several genes related to this pathway, including *PR1* are induced in different RdDM or other DNA (de)methylation mutants (Agorio and Vera, 2007; Diezma-Navas et al., 2019; Dowen et al., 2012; López et al., 2011; López Sánchez et al., 2016; Yu et al., 2013). Necrotrophic pathogens, on the other hand, are controlled by JA defense pathways, repressed in those mutants (López et al., 2011). Since our transcriptome data evidenced that SA signaling might be important for TuMV responses (Figure S2D and 2C), the general induction of SA-mediated defense pathways in the hypo-methylated mutants may be one of the mechanistic explanations of their resistance to the virus. The observed RdDM effects, however, may not be universal for plant viruses, as *ago4* mutants have been shown to be more susceptible to a tobravirus at late infection stages (Diezma-Navas et al., 2019; Ma et al., 2015). Misexpression of defense genes and changes in resistance have also been observed in histone modification mutants (Zhu et al., 2016). The genes tested here, *IBM1* and *JMJ14*, have antagonistic roles in expression regulation. In *ibm1* mutants, thousands of genes are known to gain TE-related repressive marks (Miura et al., 2009). The increased heterochromatin in this genotype therefore may possibly prevent or delay the expression of defense genes, promoting the observed susceptibility to TuMV. In contrast, JMJ14 removes H3K4 active marks from TEs and euchromatin-related marks are increased in the mutant (Lu et al., 2010; Searle et al., 2010; Yang et al., 2010). Moreover, RdDM is partially deficient in the absence of the protein (Greenberg et al., 2013). Defense genes may be therefore more primed in *jmj14* than in wild-type plants, corroborating the observed increased resistance to the virus.

As observed in other types of stresses, differences in methylation patterns in and around genes and transposons due to TuMV were observed in infected wild-type arabidopsis plants. The downregulation of some RdDM factors due to the stress (Figure 3A), together with other factors, including competition with nearby transcriptional machinery, host or viral small RNAs and viral silencing suppressor proteins, may have contributed to the observed methylation differences. Most of the DMRs were in the CpG context and mapped inside or around the transcription start site of protein-coding genes (Figure 5A, 5B and S4A). Transcription of genes having DMRs around their TSS, however, was largely not affected by the virus (Figure 6A). Absence of a significant correlation between promoter proximal CpG methylation and expression were also found in arabidopsis and other plants exposed to stress or in natural populations (Lafon-Placette et al., 2018; Mager and Ludewig, 2018; Narsai et al., 2017; Seymour and Becker, 2017; Seymour et al., 2014; Sun et al., 2019; Xu et al., 2018). Contrary to other plants, about only 5% of the arabidopsis genes are thought to be regulated by promoter methylation (Zhang et al., 2006). Furthermore, it has been shown that methylation differences in the plant are higher between tissues than in stress conditions (Seymour et al., 2014). The dilution effect produced by using whole leaves, with different cell types and likely varying viral loads, probably precluded the identification of general correlation between promoter methylation and expression. TuMV-induced genes with negative methylation and expression, though, were observed. They belonged to several functional categories and genes related to RNA or protein metabolism were the most frequent ones (Figure 6B, Table S6). Few biotic stress-related genes were found to have inverted correlations, contrary to what was observed for tobacco plants infected with cucumber mosaic virus (Wang et al., 2018). Among the identified stress-related genes, *SOMATIC EMBRYOGENESIS RECEPTOR KINASE 1* (*SERK1*) have already been linked to antiviral defense by channeling dsRNAs into PTI pathways (Niehl et al., 2016). And the HEAT SHOCK PROTEIN 22 has been associated with plant memory to cycles of heat stress (Stief et al., 2014). Those correlations should be interpreted carefully, since it is still not clear how much methylation difference is in fact required to promote significant transcriptional changes.

TEs known to be located in both euchromatic and heterochromatic environments, including centromere core, also presented differences in CHG and CHH marks, indicating a widespread deregulation of methylation machinery (Figure S4B). At 2 - 5 dpi, similar amounts of hyper- and hypo-methylation were observed in transposons, but a more pronounced hypermethylation of those elements was observed at 12 dpi (Figures 5A, 5B and S4B). Accordingly, TEs were found to be generally repressed at 12 dpi, especially against the evolved isolate (Figure 3B). This agrees with the observed repression of several TEs in arabidopsis plants infected with a DNA geminivirus (Coursey et al., 2018). Results are also in line with models predicting that early abiotic stress responses may trigger transient hypomethylation of TEs due to the overexpression of responsive genes, that can be reversed by their hypermethylation at a later time-point (Secco et al., 2015). The higher stress intensity promoted by the evolved isolate may have contributed to a faster regulation shift, explaining the increased numbers of hypermethylated and downregulated TEs at 12 dpi. Case-specific exceptions to the model were observed (Figure 3B). For example, the RdDM-targeted transposons *AtGP1* and *AtCOPIA93* were induced by TuMV infection at 12 dpi (Mirouze et al., 2009; Yu et al., 2013). A short version of the *AtCOPIA93* element, also known as *EVADÉ*, has been shown to be required for the expression of the NLR gene *RECOGNITION OF PERONOSPORA PARASITICA 4* (*RPP4*) (Zervudacki et al., 2018), a gene that was induced by both TuMV isolates (Table S1). Although the epigenetic regulation of TEs is reported to regulate expression of nearby genes, there was no clear general correlation between the state of TuMV-induced TE regulation (hyper- or hypo-methylated) and the type of nearby gene regulation (induced or repressed). This lack of correlation may reflect the dynamic changes along the infection time-course (Figure 6C), but can also be due to the small and heterochromatin-poor arabidopsis genome. In fact, DNA methylation mutants in several plants are lethal or severely compromise development, while most of them show light or no phenotype in arabidopsis (Zhang et al., 2018). There was also little correlation between TE methyl regulation and expression during the infection, in agreement with studies showing that their induction under heat stress or virus infection is not associated with DNA methylation changes (Diezma-Navas et al., 2019; Pecinka et al., 2010). Nonetheless, some DEGs that were close to TEs having non-CpG DMRs were detected, indicating a possible co-regulation mechanism. Some of them were similarly regulated by both TuMV isolates, including genes involved with disease resistance, transcriptional factors, RNA silencing and histone variants involved with stress responses. Important transcriptional and disease regulators were also found among isolate-specific cases (Table S7). As for promoter methylation differences, reported correlations should be carefully interpreted, as extra experimental approaches should be applied to confirm their causal relationships.

Apart from epigenetic factors and known RNA silencing responses, the transcriptome data also indicated that several other types of defense mechanisms were mounted against TuMV, including general shut-down of photosynthesis, metabolic rearrangements and induction of genes related to all known immunity pathways in plants (Figure 2D, S2C and S2D). The induction of several genes related to basal immunity systems based on molecular patterns (PTI) and elicitors (ETI) are in line with the increasing evidence suggesting that those types of innate defenses with conserved animal counterparts have also roles against viruses in plants (Teixeira et al., 2019). A possible role of SA in triggering defense responses was corroborated by the induction of some of its well characterized activators or targets at 12 dpi (Figure S2D, Table S1, Files S1 and S2). However, the high expression of several known SA-antagonistic genes in various time points, including WRKY25/26/33/38/62 (Kc et al., 2008; Zheng et al., 2006, 2007), indicates a possible viral counter-defense strategy. The induction of anti-SA genes was particularly high for the evolved TuMV strain (Figure S3A and Table S3). Although only two SNPs in the region coding for the viral multifunction protein VPg were detected, several lines of evidence demonstrated that the isolate had a higher virulence than the ancestral stock to arabidopsis plants. Integrating the observed methylomes and transcriptomes with virus-host protein-protein interaction networks for both isolates will be a valuable way to find the molecular basis of the adaptive process.

Viral fitness is a complex parameter often used by virologists to quantitatively describe the reproductive ability and evolutionary potential of a virus in a particular host. As cellular parasites, viruses utilize all sorts of cellular factors, reprogram gene expression patterns into their own benefit, and block and interfere with cellular defenses. All these processes take place in the host complex network of intertwined interactions and regulations. Interacting in suboptimal ways with any of such elements may have profound effects in the progression of infection and therefore in viral fitness; inefficient interactions may result in attenuated or even abortive infections. Despite its relevance, how virus evolution shapes and optimizes these interactions has received scant attention. In previous experimental evolution studies in which tobacco etch potyvirus was adapted to different ecotypes of arabidopsis, it was shown that the transcriptome of the infected plants was differentially affected depending on the degree of adaptation of the virus and identified potential host drivers of virus adaptation (Agudelo-Romero et al., 2008; Hillung et al., 2016). Here, we expand these previous studies to incorporate a new level of regulation of gene expression: DNA and histones epigenetic modifications. Our results suggest that TuMV isolates that differ in their degree of adaptation to arabidopsis may exert a differential effect on methylation patterns and in the expression of genes epigenetically regulated.

## Supporting information

All supplementary files, figures and spreadsheets

## ACKNOWLEDGMENTS

We thank Francisca de la Iglesia for technical assistance and for performing the TuMV evolution experiments in Valencia and Pawel Baster and James Barlow for technical support in Cambridge. We are grateful to Dr. César Llave and Dr. Virginia Ruiz-Ferrer for providing the *ago4, rdr2, drm1*, and *drm2* seeds and to Dr. R. Keith Slotkin for the *ddm1* ones. This work was supported by Spain Agencia Estatal de Investigación - FEDER grants BFU2015-65037-P (S.F.E.) and AGL2016-79825-R (G.G.) and by Generalitat Valenciana grant PROMETEU/2019/012 (S.F.E). R.L.C was supported by a fellowship from the Brazilian funding agency CNPq (Conselho Nacional de Desenvolvimento Científico e Tecnológico Brasil) for the stay in Valencia and from EMBO/EuropaBio for the stay in Cambridge.

## Author Contributions

R.L.C., G.G., D.C.B., and S.F.E. conceived and designed the experiments; R.L.C. and Z.K. performed infection experiments; R.L.C., A.S.C. and S.Y.M. processed and analyzed the transcriptome and methylome data; R.L.C. and S.F.E. performed statistical analysis; R.L.C., A.S.C., S.Y.M., S.L.G., G.G., D.C.B., and S.F.E. analyzed and interpreted the data; G.G., D.C.B. and S.F.E. contributed with reagents/materials/analysis tools; R.L.C. and S.F.E. wrote the paper.

## Star Methods

### Contact for Reagent and Resource Sharing

Further information and requests for resources and reagents should be directed to and will be fulfilled by the Lead Contact, Régis L. Corrêa.

### Experimental Model and Subject Details

#### Plant genotypes

For the experimental evolution and infection time-course analysis (all mRNA-seq and WGBS-seq data), wild-type *Arabidopsis thaliana* L. plants from the Col-0 ecotype were grown on short day conditions, *i*.*e*., with 8 h of light at 25 °C and 16 h in the dark at 20 °C.

Mutant arabidopsis genotypes were maintained and infected on long day conditions, *i*.*e*., with 16 h of light at 24 °C and 8 h in the dark at 20 °C. The lines *nrpD1a-3* (SALK_128428), *nrpE1* (SALK_017795C), *ibm1-4* (SALK_035608C), and *jmj14* (SALK_135712C) were obtained from the Nottingham Arabidopsis Stock Centre (NASC). Lines *rdr2-1* (SAIL_1277_H08), *drm1-2 drm2-2* and *ago4* were kindly provided by Dr. César Llave and *ddm1-2* by Dr. Keith Slotkin. Oligonucleotides used for genotyping are listed in Table S8. Primers for *ddm1-2* and *nrpD1a-3* were described elsewhere (Herr et al., 2005; Yadegari et al., 2000). All mutant genotypes were in the Col-0 background.

#### Virus isolates

The TuMV isolate YC5 (GenBank, AF530055.2) cloned under the 35S promoter and NOS terminator originally obtained from calla lily (*Zantedeschia* sp.) was used as source of virus inoculum (Chen et al., 2003). The virus was maintained in *Nicotiana benthamiana* Domin plants before being inoculated into arabidopsis plants.

### Method Details

#### Evolution experiments

Initial inoculum came from *N. benthamiana* leaf tissues infected with the YC5 TuMV isolate. Infected leaves were ground in liquid nitrogen and 100 mg of fine powder mixed with a solution containing 50 mM phosphate buffer (pH 7), 3% polyethylene glycol 6000 and 10% Carborundum at 100 mg/ml (diluted in the same PEG/phosphate buffer). Two leaves of 20 arabidopsis plants (5 weeks old) were mechanically inoculated with 5 μL of the sap. Arabidopsis plants having clear TuMV symptoms at 10 dpi were pooled and used as source of inocula as described before. A total of 10 passages of this kind were performed.

Survival analysis was done with the survfit function from the survival R package to compute Kaplan-Meier estimates. Time of “death” was scored when plants were having strong leaf yellowing symptoms (Figure S1A). Plots were generated with the survminer R package. R version 3.4.4 in RStudio was used for these analyses.

#### Infection experiments in wild-type and mutant plants

For the time-course experiments (used for all mRNA-seq and WGBS-seq data), batches of Col-0 plants were mechanically inoculated with two inocula sources: coming from benthamiana (as above) and the 9^th^ passage-infected arabidopsis tissues (ancestral and evolved TuMV, respectively). To ensure that even viral loads were used, concentration of viral transcripts in both inocula were measured by standard curve RT-qPCR. Total RNA from health and TuMV-infected arabidopsis and benthamiana plants were extracted using Plant Isolation Mini Kit (Agilent). Standard curves were constructed from eight serial dilutions of in vitro-transcribed TuMV RNA using the mMESSAGE mMACHINE^®^ SP6 Transcription Kit (Ambion). Each of the 5-fold dilutions were done by mixing viral transcripts with total RNA extracted from health tissues of arabidopsis or benthamiana for taking any PCR inhibitors into account. The 20 μL reactions were performed in an ABI StepOnePlus real-time PCR system (Applied Biosystems), using the GoTaq 1-Step RT-qPCR system (Promega). Cycling conditions were as follows: one cycle of 42 °C for 5 min and 95 °C for 10 min; 40 cycles of 95 °C for 5 min and 60 °C for 34 s; and one cycle of 60 °C for 1 min, followed by a melting curve from 60 °C to 95 °C, with 0.3 °C increments. Primers TuMV F117_F and TuMV F118_R used to amplify the viral capsid coding region are described in Table S8. Results for arabidopsis and benthamiana were analyzed separately, with their corresponding viral serial dilutions. Inoculations were performed as described above, but using adjusted tissue amounts from each plant source in order to have even inocula. Non-inoculated leaves of mock, ancestral or evolved-infected plants were collected at 2, 5 and 12 dpi and kept frozen at −70 °C until nucleic acid extraction. Plants collected at 2 and 5 dpi were symptomless. To be sure that the inoculation worked, they were left alive until the end of the time-course after leave sampling. Only frozen tissues from plants showing clear symptoms at later stages of infection were further analyzed.

Arabidopsis mutant lines were grown on long day conditions, as described above. Three-week old plants were infected with adjusted amount of TuMV-infected benthamiana or arabidopsis (9^th^ passage) saps, as described for wild-type plants. Individual plants were scored daily for typical TuMV symptoms.

#### Nucleic acid extraction and library preparation

Total DNA and RNA from TuMV-infected and healthy Col-0 plants were co-extracted using the protocol described in (Oliveira et al., 2015), with two phenol-chloroform extractions before lithium precipitation. The quality of the RNAs used for preparing mRNA-seq libraries were checked with the Bioanalyzer nano kit and quantified with the Qubit RNA BR Assay Kit (ThermoFisher). Libraries were prepared with the True-seq Stranded mRNA prep kit (Illumina), using 1 μg of total RNA as input. In total, 24 libraries were prepared, containing three biological replicates for each of the conditions. Each biological replicate was made by total RNA from individual plant systemic leaves. Libraries were sequenced with the Illumina High Output Kit v2 (2 × 75 bp) in a NextSeq 500 benchtop machine (Illumina).

DNAs (100 ng) were bisulfite-treated with the EZ DNA Methylation Gold kit (Zymo Research), before library preparation with the TruSeq DNA Methylation Kit (Illumina). In total, 18 libraries were prepared, containing two biological replicates for each condition. Each biological replicate was made by a pool of DNAs extracted from systemic leaves of three plants. Libraries were sequenced with the High Output Kit v2 (1 × 75 bp) in a NextSeq 550 benchtop machine (Illumina).

#### mRNA-seq analysis

The quality of the mRNA-seq libraries was checked with FastQC v0.11.7 (https://github.com/s-andrews/FastQC) and trimmed with TrimGalore v0.4.4 (https://github.com/FelixKrueger/TrimGalore), using cutadapt v1.3 (Martin, 2011). Twelve bases from the 5’ end of reads 1 and 2 were removed before mapping with HiSat2 v2.1.0 (Pertea et al., 2016) to the Ensembl release 39 of the *A. thaliana* TAIR10 genome assembly. For viral genome SNP calling, trimmed reads were mapped with HiSat2 to the TuMV isolate YC5 (GenBank, AF530055.2) with a modified minimum score parameter (*L*, 0-0.8) to allow more mismatches. Resulting SAM files were BAM-converted, sorted, indexed and analyzed with SAMtools v1.9 (Li et al., 2009). SNP calling was performed using bcftools v1.6 by first using the “mpileup” subroutine (with default parameters apart from -d10000) followed by the “call” subroutine as well as the “filter” subroutine filtering out low quality calls (<10). Read counting in features was done with htseq-count, using The Arabidopsis Reference Transcript Dataset (AtRTD2) (Zhang et al., 2017) as input annotation file. Differential expression analysis was done with DESeq2 v1.18.1 (Love et al., 2014), considering only genes having a total of at least 10 reads for each pairwise comparison. Functional characterization of DEGs was done with plant GOSilm implemented in the Cytoscape plugin Bingo (Maere et al., 2005) and MapMan (Thimm et al., 2004). For the analysis of differentially expressed transposons, the TEtranscripts tool was used (Jin et al., 2015). Trimmed reads were mapped with STAR (Dobin et al., 2013) to the Ensembl release 39 of the *A. thaliana* TAIR10 genome assembly. The arabidopsis transposon annotation file from TEtranscripts (http://labshare.cshl.edu/shares/mhammelllab/www-data/TEToolkit/TE_GTF/TAIR10_TE.gtf.gz) was used as input to the program.

#### RT-qPCRs

For RT-qPCRs, 1 μg of Turbo DNAse (ThermoFisher)-treated total RNAs were reverse-transcribed with Superscript IV (ThermoFisher) with random hexamer primers and used for amplification in a 10 μL reaction with the Luna^®^ Universal qPCR Master Mix (New England Biolabs). Oligonucleotides used are listed in Table S8. Amplifications were done in a CFX96 machine (Bio-Rad) with the following cycling conditions: one cycle of 95 °C for 1 min; 40 cycles of 95 °C for 15 s and 60 °C for 30 s; and one cycle of 60 °C for 1 min, followed by a melting curve from 60 °C to 95 °C. Reaction efficiencies and the fractional cycle number at threshold were calculated based on raw fluorescence with the Miner tool (Zhao and Fernald, 2005). Transcripts were quantified by the comparative ΔΔ*C*_*T*_ method, and previously known arabidopsis stable genes *PROTEIN PHOSPHATASE 2A SUBUNIT A3* (AT1G13320) and SAND (AT2G28390) were used as endogenous references (Czechowski et al., 2005). Primer sequences are described in Table S8. Primers for *ROS1* amplification were described elsewhere (Lei et al., 2015).

Relative gene expression data were fitted to generalized linear mixed models (GLMM) using plant genotypes and viral inocula as orthogonal factors. A Gamma probability distribution and a logarithm-link function were chosen based on the minimum Bayes information criterion. For testing differences among specific samples, *post hoc* pairwise comparisons with sequential Bonferroni correction tests were used. These analyses were performed with SPSS version 26 (IBM Corp.).

#### WGBS-seq analysis

The quality of the WGBS libraries was checked with FastQC v0.11.7 (https://github.com/s-andrews/FastQC) and trimmed with cutadapt v1.16 (Martin, 2011). The first nine initial and two last bases from reads were removed, and remaining ends with qscore lower than 30 were also trimmed. Reads having less than 20 bases after trimming were also discarded. Mapping was performed with Bismark - Bisulfite Mapper v0.20.0 (Krueger and Andrews, 2011), using the Ensembl release 39 of the TAIR10 genome assembly. Removal of PCR duplicates, sorting and indexing of the resulting BAM files was done with SAMtools v1.9 (Li et al., 2009). Methylation call extraction and differential analysis were performed with the Methylkit R package v1.4.1 (Akalin et al., 2012). For each pairwise comparisons (mock *vs* ancestral TuMV and mock *vs* evolved TuMV, for each time-point), bases with low (below 10×) and more than 99.9^th^ percentile of coverage in each sample were discarded before mean read normalization. Only bases covered in all samples from each pairwise comparisons were further analyzed. Methylation difference was tested with logistic regression and *P*-values were adjusted to *q*-values with the SLIM method. Differentially methylated regions in 100 bp tiles having *q* < 0.05 and methylation difference larger than 15% were selected. Assignment of each DMR to features was done with the GenomicFeatures v1.30.3 package (Lawrence et al., 2013). Annotation files were obtained from AtRTD2 (Zhang et al., 2017) and TEtranscripts tool (Jin et al., 2015). Bisulfite non-conversion rates were calculated by mapping reads to arabidopsis cytoplasmic genomes.

### Quantification and Statistical Analysis

#### General statistical analysis

Specific statistical tests used for each experiment were detailed in Figure Legends and described in the Method Details section of the Star Methods as needed.

### Data and Software Availability

The mRNA-seq and WGBS-seq data have been deposited to the SRA database under ID codes PRJNA545306 and PRJNA545300, respectively.

## Supplemental Information titles and legends

**Figure S1. Biological and molecular differences between the ancestral and evolved TuMV isolates**

(A) Categories of observed symptoms from 11 to 13 dpi. (B) Estimation by RT-qPCR of TuMV accumulation along the infection time-course. Student’s two-samples t-tests, ***P < 0.001; \*\**P* < 0.01; NS., not significant. (C) Kaplan-Meier survival regression analysis of TuMV-infected wild-type plants. Analysis based on the time each plant took to reach strong symptoms. (D) Predicted structures of the ancestral (blue) and evolved (red) VPg proteins. Altered regions were highlighted in grey. (E) Amino acid sequence alignment between the predicted VPg regions of the ancestral and evolved TuMV isolates. Shared residues are highlighted by red dots.

**Figure S2. Transcriptome responses to TuMV**

(A) Number of mapped reads to the plant or virus genomes at 2, 5 and 12 dpi. (B) Upset plot with numbers of shared DEGs in each condition. (C) Gene ontology analysis (plant GOSlim) for the identified DEGs. Circle size represents level of enrichment. (D) Visualization of cluster 40 of the AtCKN Arabidopsis network in response to the evolved TuMV at 2, 5 and 12 dpi.

**Figure S3. Transcriptional profiles of biotic- and abiotic-related genes**

(A) Transcriptional profiles of transcription factor genes. (B) RT-qPCR confirmation of biotic, abiotic and development genes in TuMV-infected plants at 12 dpi. Relative gene expression data were fitted to a generalized linear mixed model (GLMM) with plant genotype (*PR1, WAK1, HSP70*, and *COR15a*) and source of inocula (mock, ancestral and evolved viruses) incorporated as orthogonal factors. (C) RT-qPCR confirmation of *IBM1* and *ROS1* genes in TuMV-infected plants at 12 dpi. Relative gene expression data were fitted to a GLMM with plant genotype (*IBM1* and *ROS1*) and source of inocula (mock, ancestral and evolved viruses) incorporated as orthogonal factors. (B) and (C) a Gamma probability distribution and a logarithm-link function were assumed. *Post hoc* pairwise comparisons with sequential Bonferroni correction tests were performed; \*\*\**P* < 0.001; \*\**P* < 0.05; \**P* < 0.1; NS., not significant.

**Figure S4. Whole-genome bisulfite sequencing (WGBS) of infected wild-type plants**

Number of hyper- or hypo-methylated DMRs in the three cytosine contexts (CpG, CHG and CHH) are presented for each condition. (A) Total number of DMRs found in the genome. (C) Number of DMRs in selected TE families.

**Table S1. List of all DEGs identified in the DESeq2 analysis (adjusted *P* < 0**.**05) for each TuMV infection condition**.

**Table S2. List of DEGs regulated by all TuMV infection conditions**. Related to Figure S2B.

**Table S3. List of selected DEGs regulated by TuMV infection**. Related to Figures 3A, S3A, S3B and S3D.

**Table S4. List of differentially expressed transposons obtained with TEtranscripts (adjusted *P* < 0**.**05) for each TuMV infection condition**.

**Table S5. Genomic ranges and values for all identified DMRs in each TuMV infection condition**.

**Table S6. Genes having negative correlation between transcriptional start site proximal methylation and expression**. Related to Figures 6A and 6B.

**Table S7. DEGs close to TEs with non-CpG DMRs at 12 dpi**. Related to Figure 6C.

**Table S8. List of primers used in this study**.

**File S1. Dynamic visualization of cluster 40 of the AtCKN Arabidopsis network in response to the ancestral TuMV isolate at 5 and 12 dpi**.

**File S2. Dynamic visualization of cluster 40 of the AtCKN Arabidopsis network in response to the evolved TuMV isolate at 2, 5 and 12 dpi**.

