## Supplementary figures and images for "Viral fitness determines the magnitude of transcriptomic and epigenomic reprogramming of defense responses in plants"

### FigS1.png

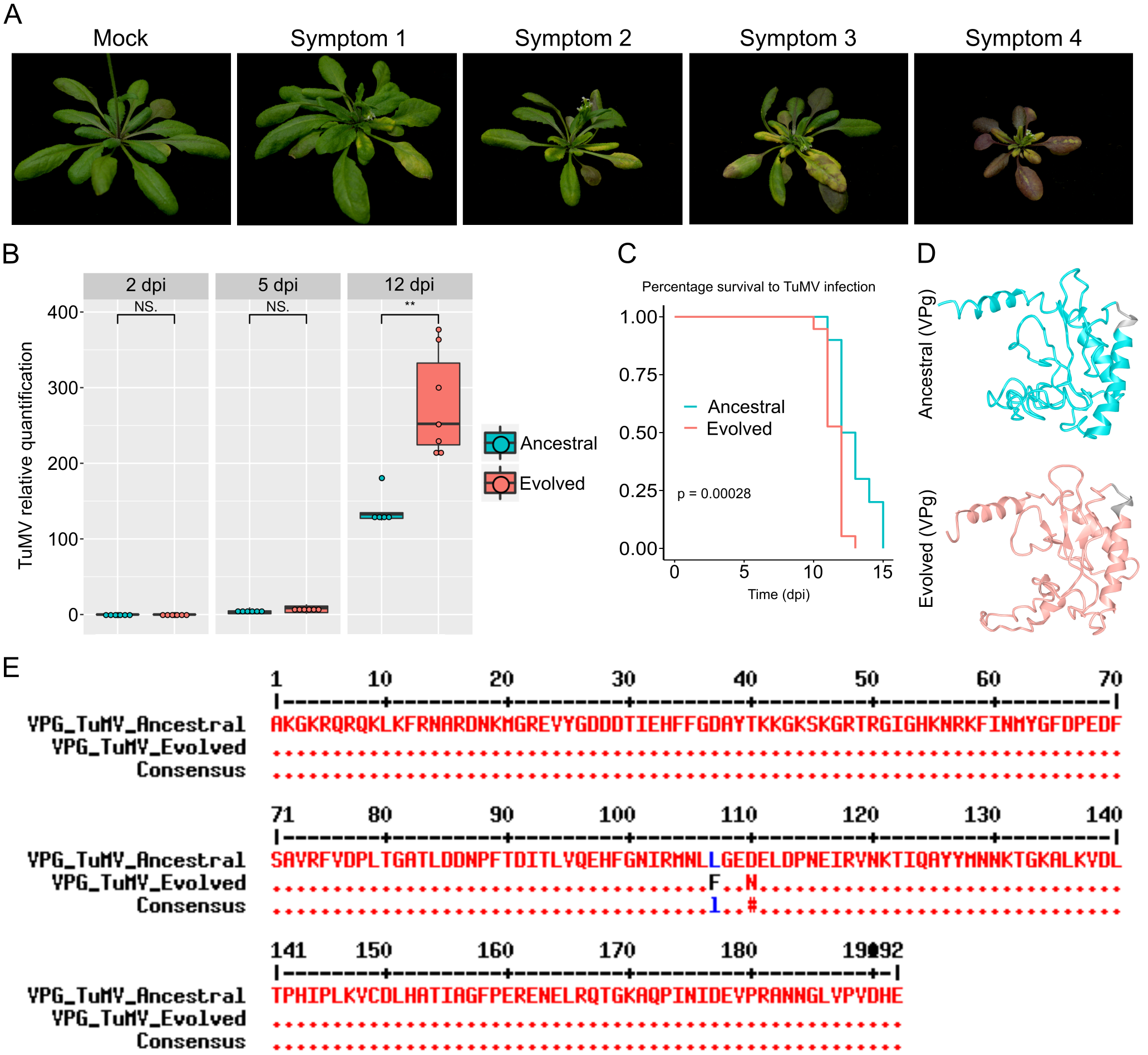

### FigS2.png

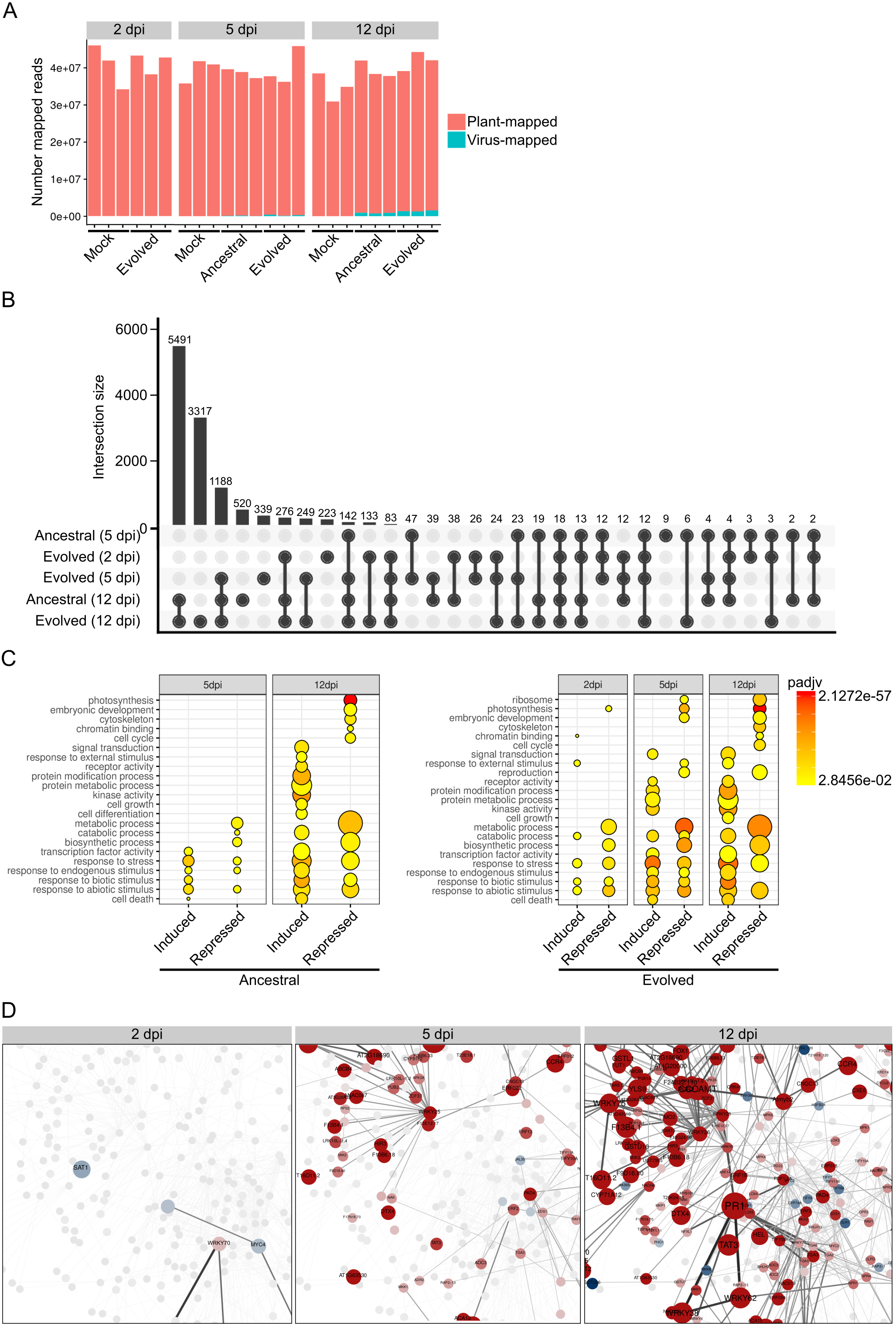

### FigS3.png

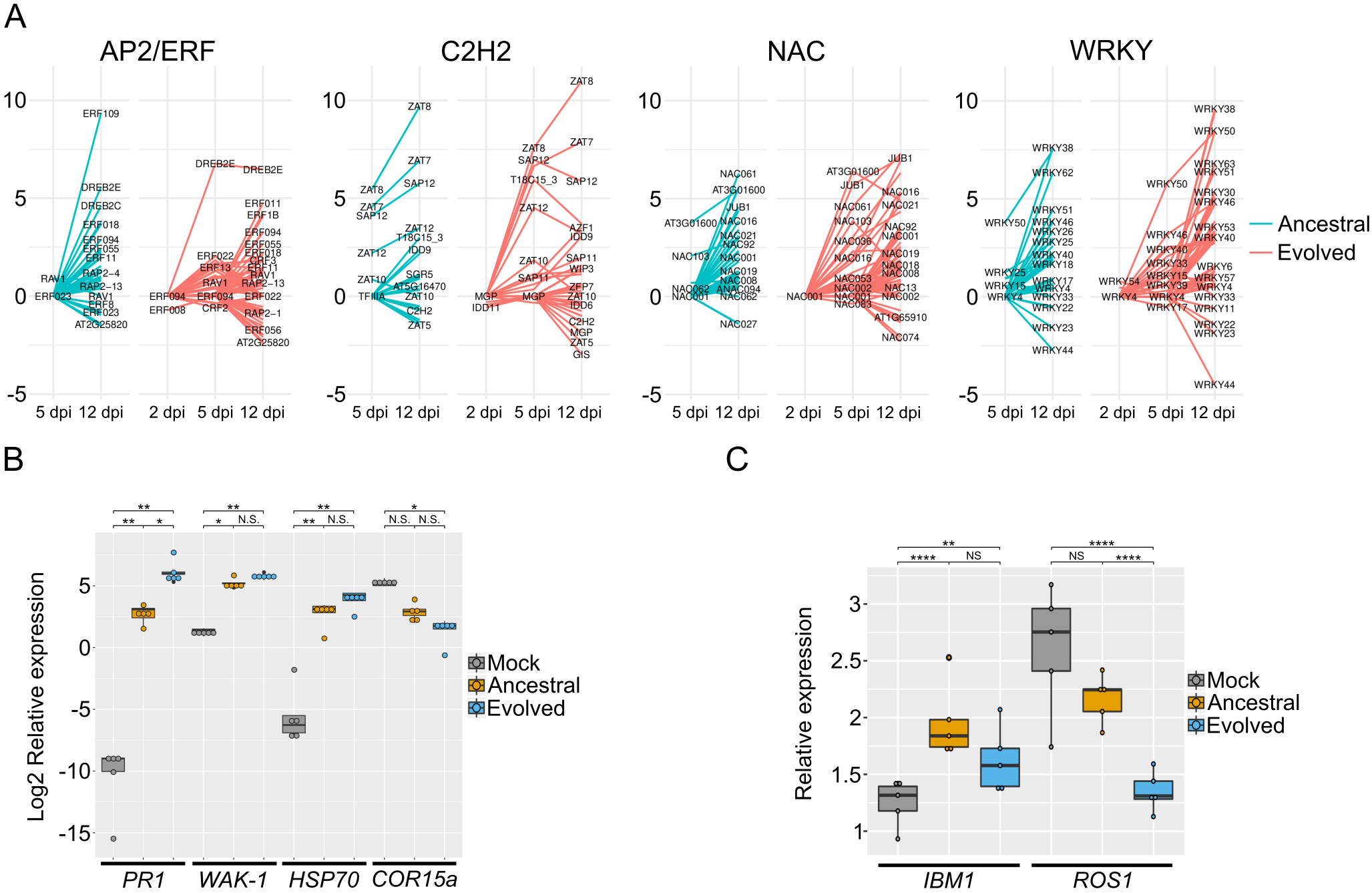

### FigS4.png

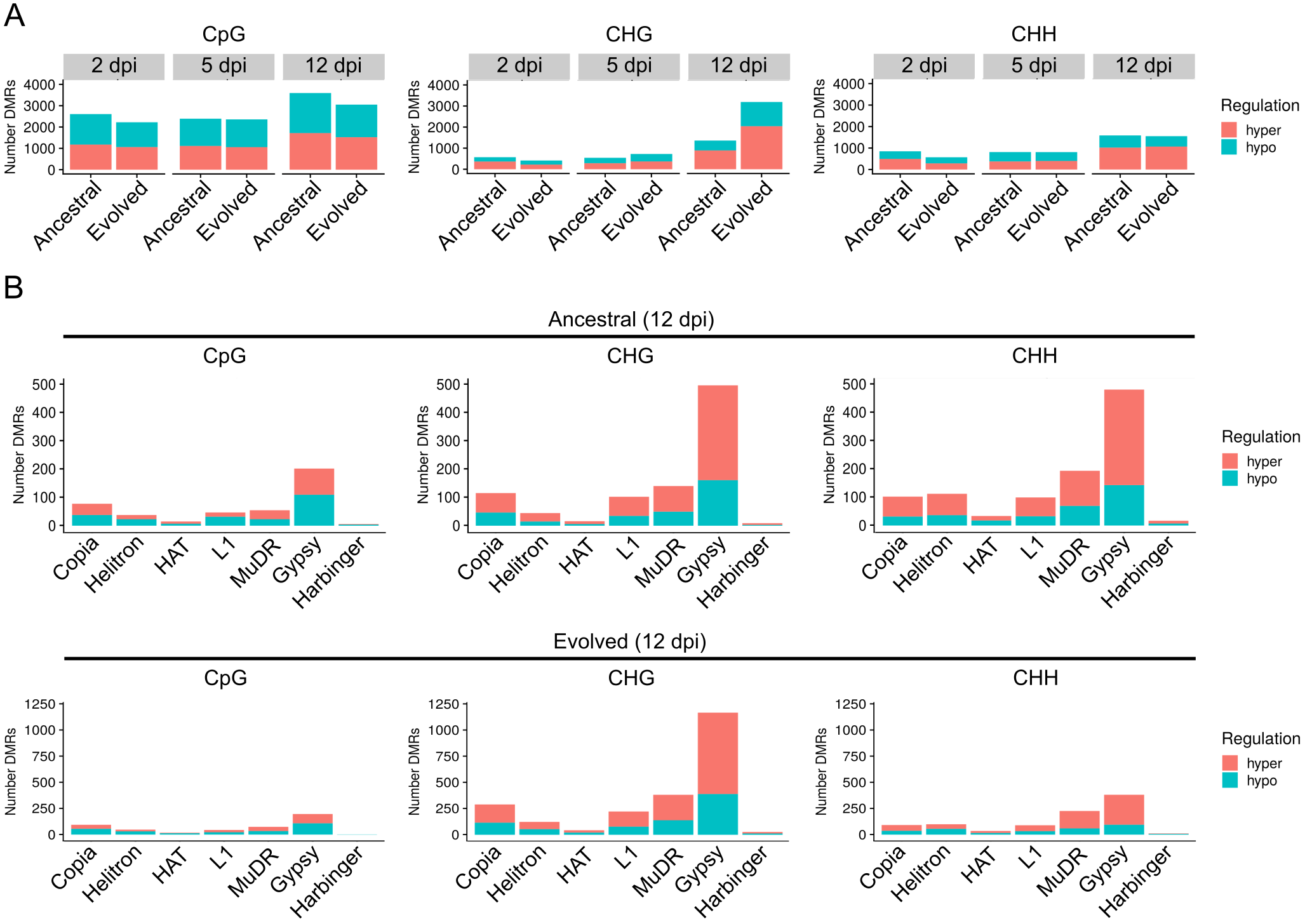
